## Supplementary material for "A pipeline for constructing reference genomes for large cohort-specific metagenome compression": Supplymentary Figures

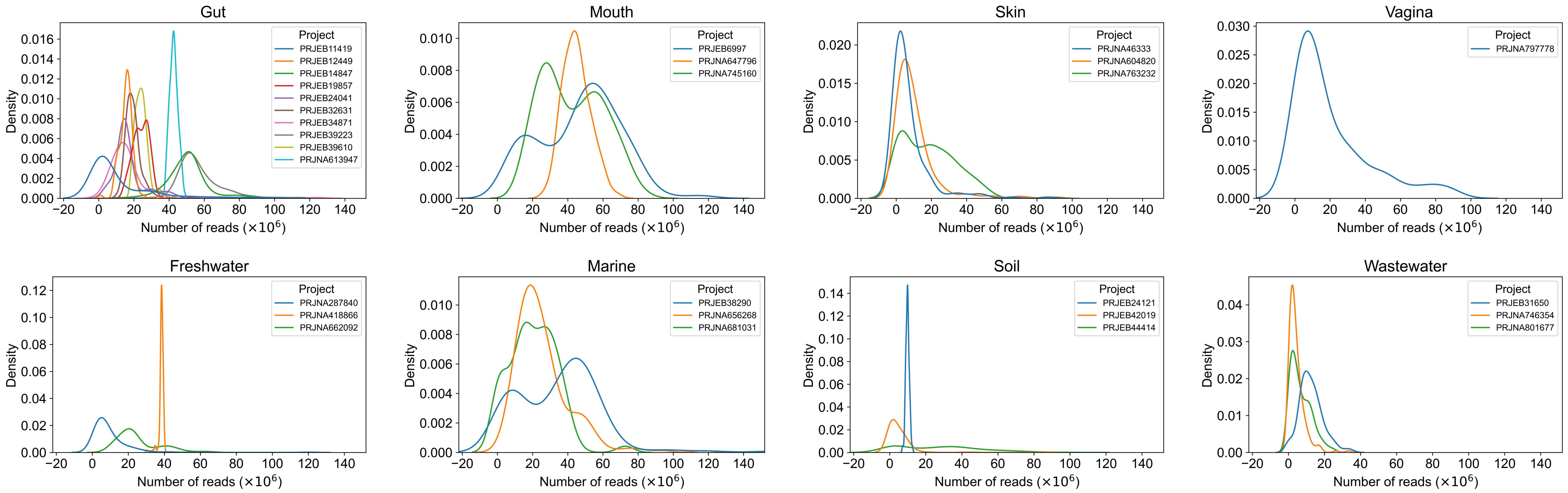


**Figure S1: Kernel distribution estimation plot of sequence depths within each dataset used in this study.**

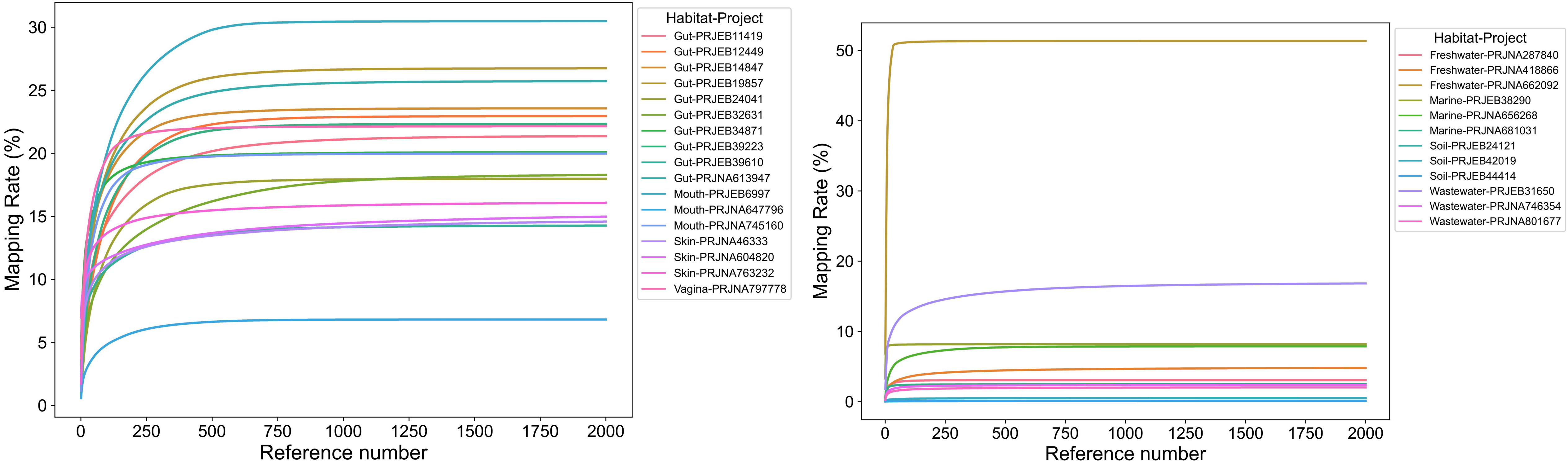


**Figure S2: Variation of the mapping rates of various datasets with the number of genomes.** Sequencing data of 50 samples from each dataset were aligned with the basic reference database (MetaPhlAn3_db_25k for human-associated datasets and MetaPhlAn3_db_25k + Env_db_6k for environment-associated datasets). Reference genomes in the database were sorted in descending order based on the number of reads mapped, and mapping rate is calculated for the top 1-2000 reference genomes. Mapping rate = Number of reads mapped to these reference genomes / total reads of all samples.
